# Petrobactin protects against oxidative stress and enhances sporulation efficiency in *Bacillus anthracis* Sterne

**DOI:** 10.1101/350611

**Authors:** Ada K. Hagan, Yael M. Plotnick, Ryan E. Dingle, Zachary I. Mendel, Stephen R. Cendrowski, David H. Sherman, Ashootosh Tripathi, Philip C. Hanna

## Abstract

*Bacillus anthracis* is a gram-positive bacillus that under conditions of environmental stress, such as low nutrients, can convert from a vegetative bacillus to a highly durable spore that enables long-term survival. The sporulation process is regulated by a sequential cascade of dedicated transcription factors but requires key nutrients to complete, one of which is iron. Iron acquisition by the iron-scavenging siderophore petrobactin is the only such system known to be required for vegetative growth of *B. anthracis* in iron-depleted conditions, *e.g.*, in the host. However, the extent to which petrobactin is involved in spore formation is unknown. This work shows that efficient *in vitro* sporulation of *B. anthracis* requires petrobactin, that the petrobactin biosynthesis operon (*asbA-F*) is induced prior to sporulation, and that petrobactin itself is associated with spores. Petrobactin is also required for both oxidative stress protection during late stage growth and wild-type levels of sporulation in sporulation medium. When considered with the petrobactin-dependent sporulation in bovine blood also described in this work, these effects on *in vitro* growth and sporulation suggest that petrobactin is required for *B. anthracis* transmission via the spore during natural infections in addition to its key functions during active anthrax infections.

**Importance:** *Bacillus anthracis* causes the disease anthrax, which is transmitted via its dormant, spore phase. However, converting from bacilli to spore is a complex, energetically costly process that requires many nutrients including iron. *B. anthracis* requires the siderophore petrobactin to scavenge iron from host environments. We show that in the Sterne strain, petrobactin is required also for efficient sporulation, even when ample iron is available. The petrobactin biosynthesis operon is expressed during sporulation, and petrobactin is biosynthesized during growth in high iron sporulation medium but instead of being exported, the petrobactin remains intracellular to protect against oxidative stress and improve sporulation. It is also required for full growth and sporulation in blood (bovine), an essential step for anthrax transmission between mammalian hosts.

## Introduction

*Bacillus anthracis* is a gram-positive, spore-forming bacillus, which causes the disease anthrax. In humans, anthrax can manifest in four ways depending on the route of exposure to *B. anthracis* spores: cutaneous, inhalational, gastrointestinal, or injectional (1, 2). Following aerosol exposure, the spores, a metabolically dormant form of *B. anthracis*, are taken up by antigen presenting cells (APCs) such as macrophages and dendritic cells (3, 4). While associated with APCs, a set of small molecules from the host initiate germination of spores into vegetative bacilli (3). The bacilli rapidly initiate cellular functions and within 30 minutes begin transcription and translation of required proteins, including the toxins that both enable escape from the APC and cause anthrax pathologies (5). If the APC is in transit to proximal lymph nodes when escape occurs, the bacilli are released directly into the blood or lymph to replicate, quickly reaching titers greater than 10^8^ CFU/mL (6, 7).

The *B. anthracis* spore is the infectious particle for its role in anthrax transmission, and a dormant structure enabling survival of harsh conditions including: nutrient deprivation, extreme temperatures, radiation, and desiccation (6, 8–11). As nutrients diminish and cell density increases, environmental sensors initiate a cascade of transcriptional regulators to construct a spore from both the inside out and the outside in (12–14). Most of the research describing sporulation has been conducted in *B. subtilis* and will be described in brief here (see 10 for a recent review).

The first morphological change observed during sporulation is asymmetric division of a bacillus into the mother cell and prespore compartments, which is initiated by phosphorylation of the transcriptional regulator Spo0A and activation of the sporulation-specific sigma factor σ^H^ (12, 15, 16). The next step in transcriptional regulation is the compartmentalized activation of two early sporulation sigma factors, σ^F^ and σ^E^, in the prespore and mother cell, respectively (12). A suite of σ^F^-and σ^E^-dependent proteins enable engulfment of the prespore by the mother cell in the second major morphological change (12, 17). Final maturation of the spore is regulated by the prespore-specific σ^G^ and the mother-cell-specific σ^K^ (12,18,19). When completed, the spore structure is composed of a dehydrated core, containing the genome and silent transcriptional and translational machinery, surrounded by an inner membrane; a layer of modified peptidoglycan known as the cortex; an outer membrane; a proteinaceous spore coat; and, for *B. anthracis*, the exosporium (9,12,14).

Sporulation is an energetically costly process. While sporulation is initiated by nutrient depletion, efficient sporulation still requires access to many nutrients, including large amounts of iron (1.5-2mM) (20, 21). Iron is required as a cofactor for enzymes requiring electron transfer such as those involved in environmental sensing, ATP synthesis, and the tricarboxylic acid cycle (22). To scavenge iron from the environment during low iron availability, many bacteria can synthesize small molecules called siderophores. In iron-replete conditions, however, siderophores and other iron acquisition systems are repressed by the ferric uptake repressor Fur, or a similar system. Fur is a dual iron and DNA-binding protein. In the iron-bound form, Fur tightly binds sequences known as Fur-boxes thus repressing transcription of any downstream genes. Low-iron stress causes the iron to be shunted from Fur to essential cellular processes, which de-represses Fur-regulated genes allowing for expression of iron acquisition systems (23, 24).

While iron is essential, excess free iron is toxic to the cell, which thus requires dedicated proteins to prevent the formation of superoxide radicals via participation of iron in the Fenton reaction. Iron in *B. anthracis* is sequestered by ferritins, the mini-ferritin DPS, and superoxide dismutases (25, 26). These proteins contribute to iron storage in *B. anthracis* spores (~10μM), which is presumed to be required for outgrowth from the spore in iron-limiting conditions (*e.g.*, within an APC endosome), until active iron acquisition systems can be expressed one to two hours following germination (27, 28). One such system is the siderophore petrobactin, whose biosynthetic machinery is encoded by the *asb* operon and is induced within two hours of germination (27, 29).

*B. anthracis* has three known active iron acquisition systems: two siderophores, petrobactin and bacillibactin, and a heme acquisition system. Of the three systems, only petrobactin is required for growth in macrophages and virulence in a murine inhalational anthrax model (29, 30). Previous studies have elucidated much about petrobactin use in *B. anthracis* including defining: the biosynthetic pathway for petrobactin (*asb* operon), the petrobactin-iron complex receptor (FhuA), import permeases (FpuB/FatC/FatD), ATPases (FpuC/FatE), and the petrobactin exporter (ApeX) (29, 31–34). However, previous studies have also suggested that the *asb* operon may be regulated by environmental conditions other than iron (35, 36). In the current work, we investigated whether petrobactin-dependent iron acquisition plays a role in aspects of *B. anthracis* Sterne spore biology and the associated regulation of *asb*.

## Results

### Petrobactin is required for sporulation but not germination

Spores cannot be infectious particles without first germinating to the vegetative state, so to begin evaluating the role of petrobactin in spore biology, initial experiments investigated the effect of petrobactin on germination in low-iron conditions. To observe germination kinetics, spores of wild-type *B. anthracis* Sterne, an *asb* (petrobactin-null) mutant strain, and a *dhb* (bacillibactin-null) mutant strain were incubated in iron-depleted medium supplemented with 1mM inosine (IDM+I) for one hour. The *asb* mutant did not display a defect in germination, relative to either wild-type or *dhb* mutant spores (Figure 1A).

To further explore our hypothesis that petrobactin plays a role in spore biology, we tested the ability of an *asb* mutant strain and a *dhb* mutant strain to sporulate relative to wild-type. At 12 and 36 hours of growth in sporulation medium, colony forming units per mL (CFU/mL) were enumerated to determine total and sporulated counts. Despite an abundance of ferrous iron (1.7mM) in the medium and growth to 10^7^ CFU/mL (Figure 1B), less than 10^6^ CFU/mL (~10%) of the *asb* mutant strain population had sporulated (Figure 1C). That was nearly two log fewer spores than the wild-type and *dhb* mutant strains whose spore populations exceeded 10^7^ CFU/mL at 12 hours post-inoculation (Figure 1C). This defect in sporulation by the *asb* mutant strain was not observed at 36 hours post-inoculation, suggesting the petrobactin phenotype is kinetic. As a control, the defect was rescued at 12 hours by supplementing the *asb* mutant strain with 25μM of purified petrobactin at inoculation (Figure 1B, 1C). Since sporulation of the *asb* mutant strain can be complemented *in trans* with purified petrobactin, these data suggest that petrobactin is biosynthesized and that the *asb* operon is expressed prior to spore formation in this growth condition, despite the presence of high iron levels.

**Figure 1.**
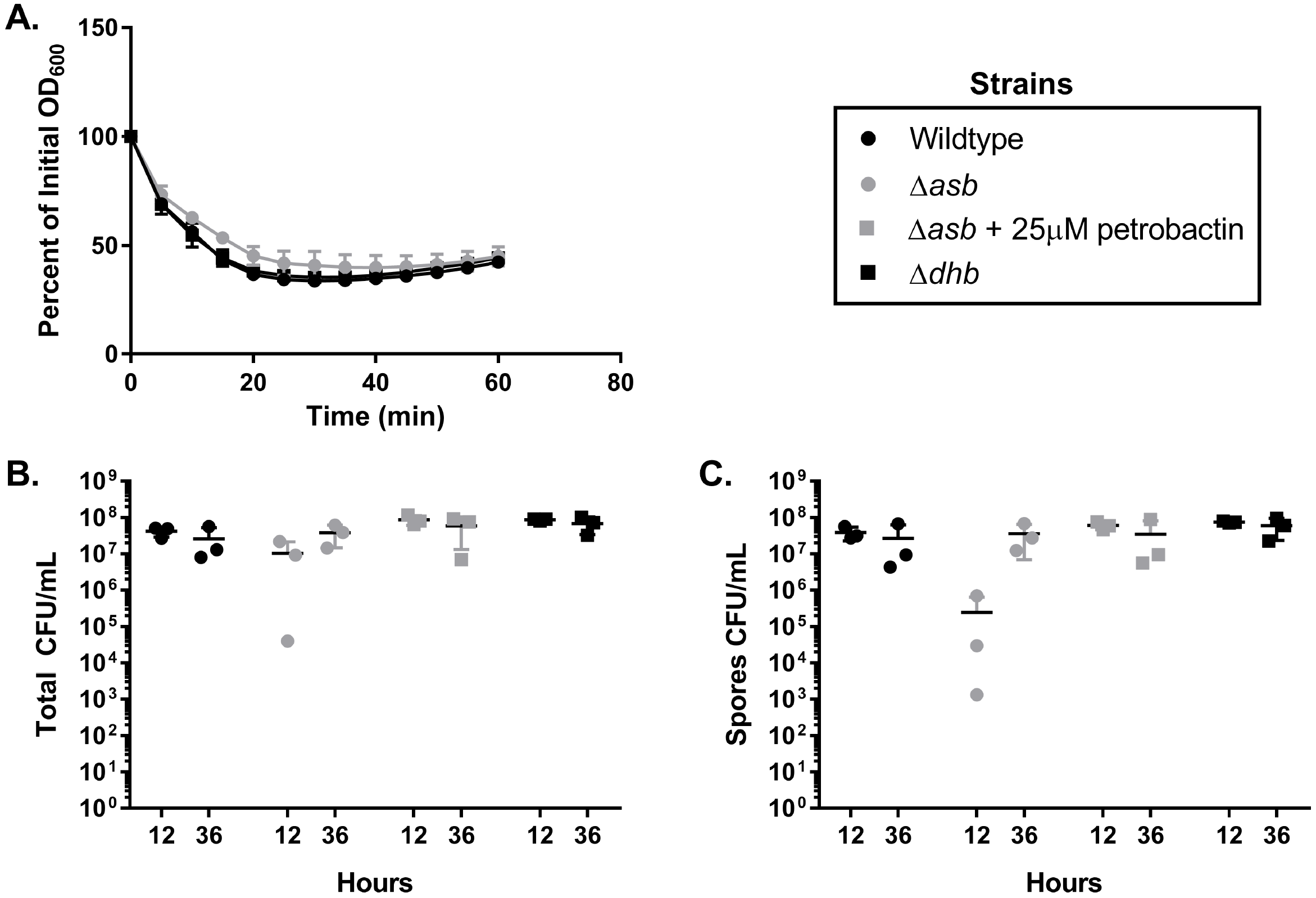
*asb* mutant spores germinate but fail to sporulate efficiently. **A)** Wild-type, *dhb* mutant, and *asb* mutant spores were inoculated in IDM + 1mM inosine at a starting OD_600_ between 0.25 and 0.5. The OD_600_ was measured every five minutes for one hour. Data are presented as percent of initial OD_600_ and are representative of n=3. **B-C)** overnight cultures of wild-type, *asb* mutant ± 25μM petrobactin, and *dhb* mutant bacilli were incubated in three mL of ModG medium and incubated at 37°C with shaking. At 12 and 36 hours post-inoculation the **B)** total and **C)** spore CFUs/mL were determined by serial dilution and plating. Data are compiled from three independent experiments.

### The *asb* operon is transcribed and translated during late stage growth and early sporulation

To understand how *asb* might be expressed despite iron levels capable of suppressing expression during vegetative growth, we used the <u>d</u>ata<u>b</u>ase of transcriptional regulation in *<u>B</u>acillus <u>s</u>ubtilis* (DBTBS) prediction tool to search the 500bp upstream of *asbA* for putative sigma factor binding sites (37). There were potential consensus binding sites for two sporulation-specific sigma factors (σ^G^ and σ^K^), the general stress transcription factor σ^B^ and the oxidative stress response regulator PerR encoded in this region, all identified with at least 95% confidence (Figure 2A). This suggests that (an) alternative regulation system(s) may be active during sporulation and could be responsible for the petrobactin-dependent sporulation phenotype. To characterize expression of the operon, we generated fluorescent reporters. Two reporter constructs for the *asb* operon, one transcriptional and one translational, were generated by fusing the 500bp upstream of *asb* and the first eight codons of *asbA* (Figure 2A, underlined)—either separated by a ribosomal binding site (transcriptional) or directly (translational)—to the green fluorescent protein allele *gfpmut3α* (38). To facilitate wildtype-like expression of the reporters, each was inserted on the *B. anthracis* genome immediately downstream of the *asbF* transcriptional terminator by allelic exchange.

To measure reporter expression, strains of the transcriptional and translational reporters, along with the wild-type strain, were grown in sporulation medium with shaking for 12 hours and the OD_600_ and GFP fluorescence measured every five minutes. The transcriptional and translational reporter strains both grew identically to the wild-type strain (Figure 2B) and the calculated RFUs indicate that *asb* is both transcribed and translated during stationary phase growth in sporulation medium (Figure 2B, black arrows).

After observing that the *asb* operon is expressed during late stage growth in sporulation medium, we next sought to determine if any of the predicted sporulation sigma factors were required for *asbA-F* expression. Here, we used plasmid-based transcriptional reporter constructs where the 260bp upstream of *asbA* (Figure 2A, vertical line) were fused to *gfpmut3α*, cloned into the pAD123 expression vector, and expressed in a wild-type *B. anthracis* Sterne background. This construct lacks the predicted binding sites for sporulation-specific sigma factors σ^G^ and σ^K^ but retains predicted binding sites for Fur, σ^B^, and PerR (Figure 2A).

To measure expression of *asbA-F* by this construct, strains of wild-type, the transcriptional reporter, and a promoter-less *gfpmut3α* were grown in sporulation medium as described and the RFUs were similarly calculated. Overall growth kinetics were similar and the 260bp *asb* promoter was sufficient for *asb* transcription during late stage growth (Figure 2C, black arrow). The observed increase in RFU for Figure 2B versus 2C is likely an artifact from increased copy numbers of plasmid-based reporters. Together these data suggest that the high iron levels in the sporulation medium do fully repress the *asb* operon by Fur and that sporulation-specific sigma factors are not required for expression of *asbA-F* during these conditions.

**Figure 2.**
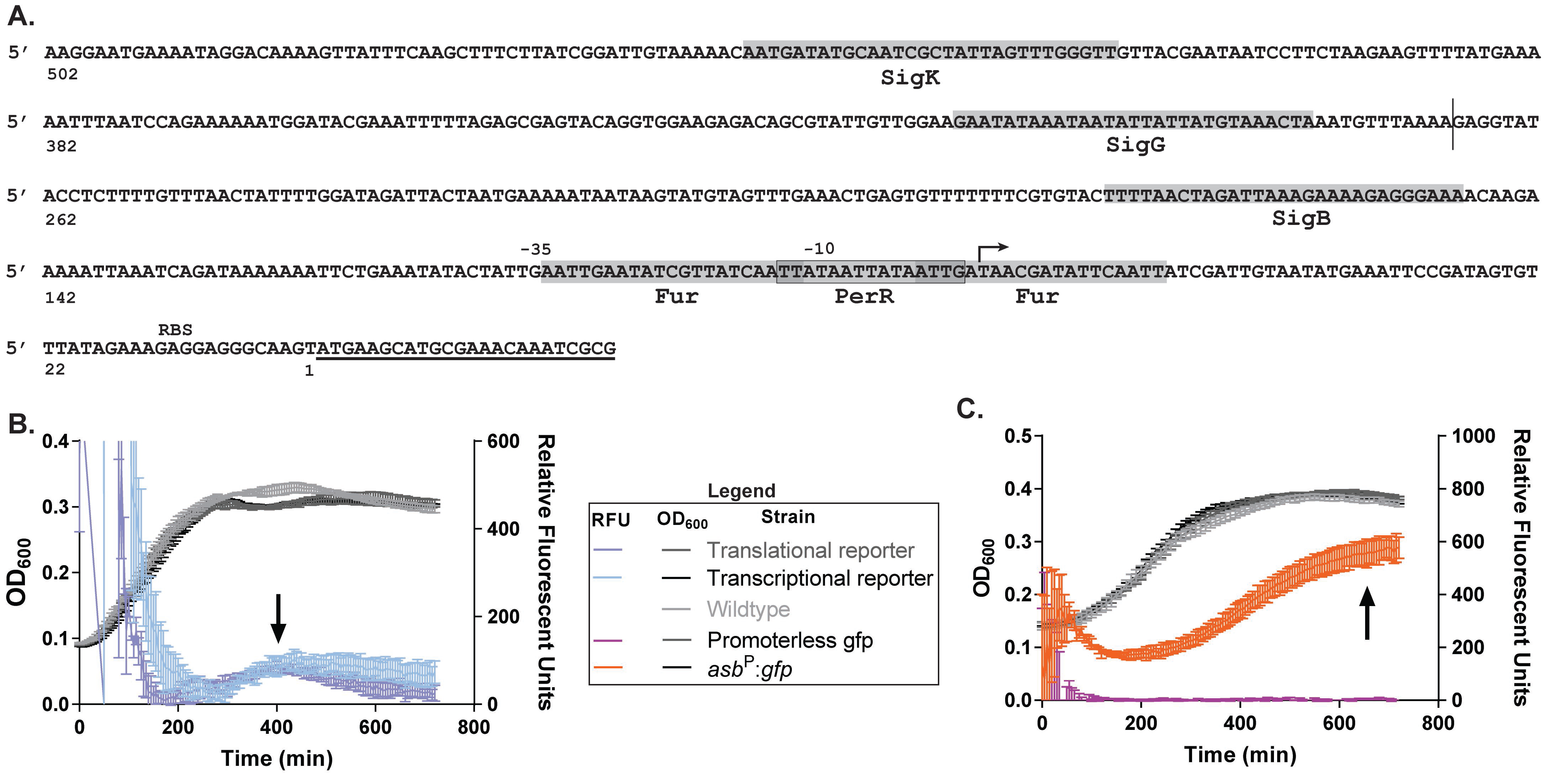
The *asb* transcriptional and translational fluorescent reporters illuminate expression during late stage growth. **A)** Schematic of putative transcriptional regulator binding sites (shaded regions) upstream of *asbA* (underlined). The bent arrow denotes the transcriptional start site in IDM (unpublished data, 34), vertical line indicates start of primer for plasmid-based reporter. **B-C)** Wild-type and fluorescent reporters were inoculated into ModG (+ 10μg/mL chloramphenicol as needed) at a starting OD_600_ of 0.05. Growth (Left axis, OD_600_) and relative fluorescence units (Right axis) were measured every five minutes for 12 hours. Data are representative of three independent experiments. Black arrows indicate late stage expression of the *asb* operon **B)** genomic-based *asb* transcription and translation *gfpmut3α* fusion reporters **C)** plasmid-based promoterless *gfpmut3α*, and *asb:gfpmut3α* transcriptional reporters.

Given these conclusions, we next wanted to better understand the population dynamics and kinetics for *asb* expression relative to sporulation. The chromosome-based translational reporter and wild-type strains grown in sporulation medium were imaged with phase-contrast and fluorescence microscopy at six, eight, ten, and 12 hours post-inoculation. Individual bacilli were scored for Gfpmut3α expression (positive is at least 1.4× above background fluorescence) and sporulation (if they contained phase bright spores) (representative images in Figure 3B-E, for wild-type see Supplementary Figure 1). At six hours of growth, 100% of the translational reporter cells were fluorescent, thus expressing the *asb* operon (Figure 3A-B). The number of fluorescent bacilli decreased over time, with 80% of the population expressing *asb* at eight hours of growth and only 20% at 10 hours of growth (Figure 3A, 3C-D). No bacteria were scored as fluorescent at the 12-hour time point (Figure 3A, 3E). Phase bright spores were not observed until ten hours post-inoculation, at which point spores were present in 65% of bacilli (Figure 3A, 3D). At 12 hours post-inoculation, 90% of the population were either sporulating or mature, free spores (Figure 3A, 3E). Together with the data from Figure 2B-C, these data indicate that *asb* expression peaks and terminates before maturation to phase-bright spores and likely before the onset of sporulation (especially given the long half-life of Gfpumut3α).

**Figure 3.**
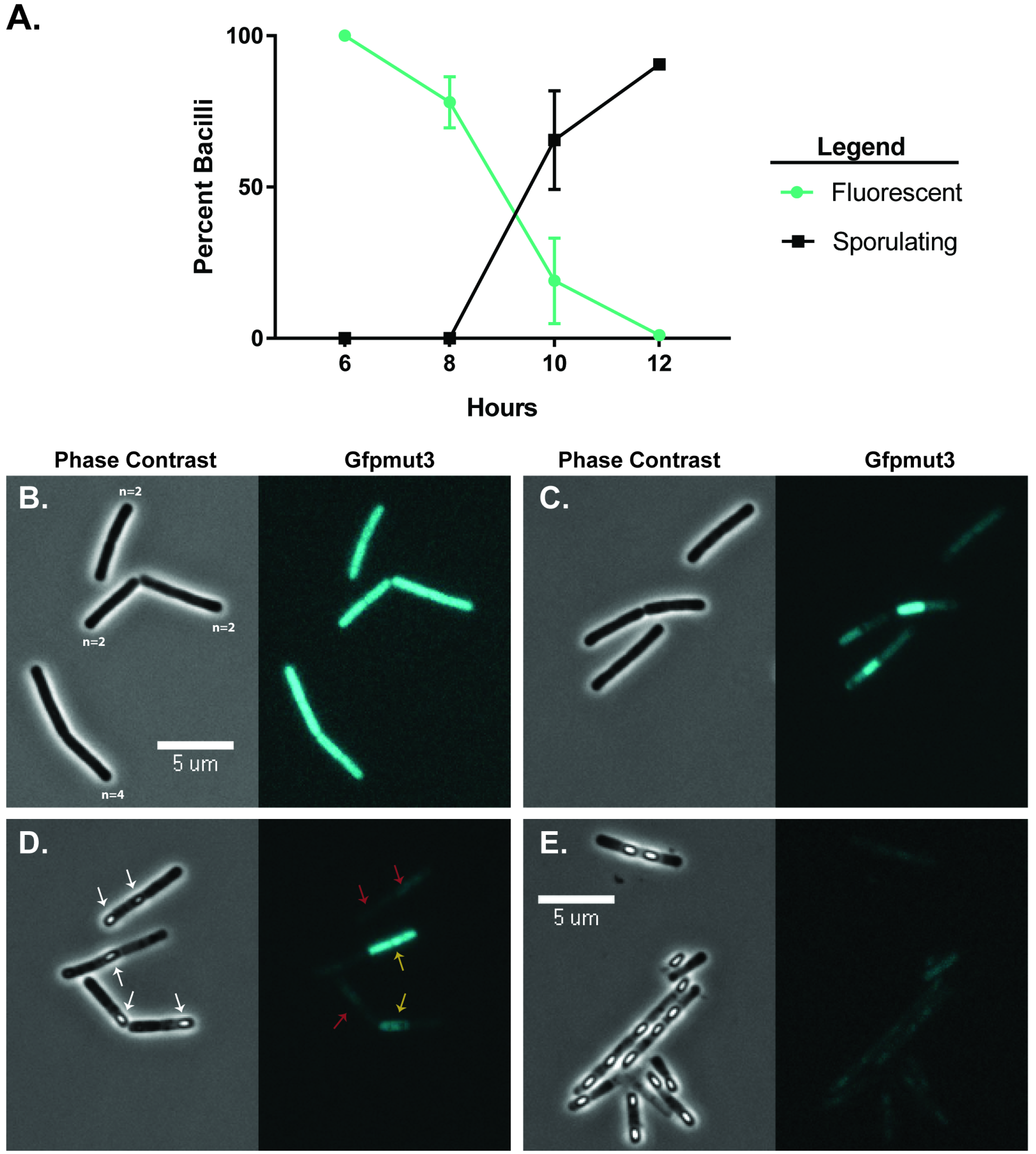
Translation of *asb* shuts down during late stage growth while sporulation occurs. The *asb* translational reporter strain was grown in ModG sporulation medium with both phase contrast and Gfpmut3α fluorescent micrographs taken at six, eight, ten, and 12 hours of growth and the bacteria scored for fluorescence and sporulation. **A)** Pooled data from two replicates of the percent of bacilli fluorescent and/or sporulating over time **B-E)** Representative phase-contrast and fluorescence images from each time point. **B)** six hours, n=X indicates the number of bacilli counted per chain. **C)** eight hours **D)** ten hours, examples of scoring for phase bright spores (white arrows), fluorescent (yellow arrows) and non-fluorescent (red arrows). **E)** 12 hours.

These observations confirm that *asb* expression does not require sporulation-specific sigma factors during sporulation (Figure 2C), particularly since both σ^G^ and σ^K^ are active during the later stages of sporulation by which point fluorescence has markedly declined, probably due to either degradation or protein dilution due to cell division. So, while petrobactin does not appear to be required for the process of sporulation *vis a vis* sporulation-specific regulation of *asb*, it is required for efficient sporulation and since cell stress precedes sporulation, *asb* expression may be induced by a stress response regulator such as σ^B^ or PerR.

### During sporulation, petrobactin is not exported, remains associated with the spore and is protective against oxidative stress

The petrobactin requirement for efficient sporulation and the upregulation of *asbA-F* during this period suggest that petrobactin is synthesized and may be present in the culture medium. However, the petrobactin-specific catechol moiety 3,4-dihydroxybenzoate, was not detected in sporulation medium at 12 hours post-inoculation by the colorimetric catechol assay (data not shown). This could be due to either assay interference by the medium, petrobactin levels below the limit of detection, or suggest an intracellular role for petrobactin. To confirm petrobactin biosynthesis and address these possibilities, we used laser ablation electron spray ionization mass spectroscopy (LAESI-MS) to detect petrobactin both in the spent culture medium and the cell pellets of *B. anthracis* wild-type and *asb* mutant strains grown in sporulation medium for 12 hours (33). When compared against our negative control, the petrobactin-null *asb* mutant strain, LAESI-MS confirmed the catechol assay results as it did not detect petrobactin in the spent culture medium from the wild-type strain (Figure 4A), indicating no discernable export of this siderophore took place. However, petrobactin was detected in cells of the wild-type strain thus confirming synthesis (Figure 4A).

The use of petrobactin intracellularly might result in association of petrobactin with the *B. anthracis* Sterne spore so we also subjected wild-type and *asb* mutant strain spores to LAESI-MS (n=3) analysis. This experiment detected petrobactin in wild-type, but not *asb* mutant strain spores (Figure 4B). This phenotype could be restored by supplementing growth and sporulation of the *asb* mutant strain with 25μM of purified petrobactin (n=1). Complete ablation of the spores was confirmed by an abundance of the spore-core-component calcium dipicolinic acid in the chromatograph (data not shown). These data indicate that while petrobactin is not exported into the medium at detectable levels, it is biosynthesized but remains associated with the spore.

**Figure 4.**
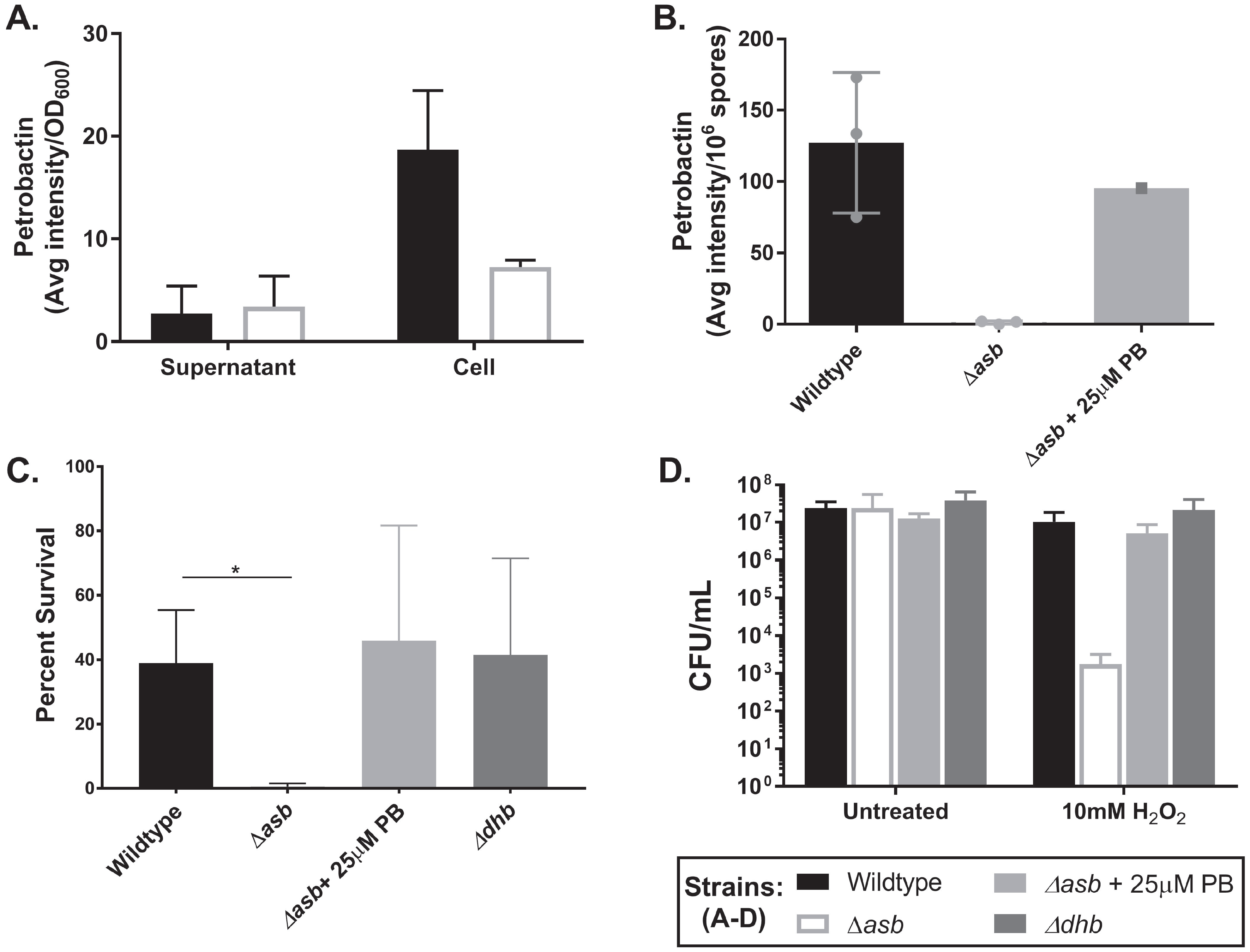
Petrobactin has an intracellular role to protect against oxidative stress and associates with the *B. anthracis* spore. **A)** LAESI-MS analysis of petrobactin in the supernatants and cell pellets of wild-type and *asb* mutant strains grown in sporulation medium for 12 hours. Data are presented as counts per OD_600_ and pooled from three independent experiments. **B)** LAESI-MS analysis of petrobactin in 6×10^7^ wild-type, and *asb* mutant ± 25μM petrobactin (n=1) spores harvested from ModG medium. **C-D)** Oxidative stress survival by wild-type, the *asb* mutant strain ± 25μM petrobactin, and the *dhb* mutant strain grown in ModG medium for eight hours following incubation with either water (untreated) or 10mM H_2_O_2_ for ten minutes at 37°C. Samples were serially diluted and plated to calculate **C)** percent survival from **D)** CFU/mL. Data are pooled from three independent experiments and analyzed using an unpaired t-test *p-value < 0.05.

To this point in our studies, the mechanism of *asb* expression and role for petrobactin biosynthesis in sporulation remains unclear. Binding sites for *asbA-F* regulators in the plasmid-based *asb* transcriptional reporter include PerR, an oxidative stress response regulator, and σ^B^, a general stress response regulator (Figure 2A). In *B. subtilis*, σ^B^ is active during early sporulation, but is not required for either sporulation or an oxidative stress response, likely since most σ^B^-regulated genes can be activated by other transcription factors (40, 41). However, Lee et al., found that oxidative stress can induce petrobactin expression and synthesis, even in high iron conditions (35). While sporulation is not known to be preceded by oxidative stress, *B. subtilis* cells become resistant to oxidative stress upon entry to the stationary phase (41–43).

Additionally, oxidative stress protective enzymes are induced during late stage growth of *B. anthracis*, maintained during sporulation, and two superoxide dismutases become incorporated in the exosporium (27, 28). Taken together with evidence of intracellular petrobactin, we predicted that petrobactin is protective against oxidative stress.

To test this hypothesis, wild-type, *asb* mutant ± 25μM petrobactin, and *dhb* mutant strains were tested for resistance to the oxidative stressor hydrogen peroxide (H_2_O_2_) at eight hours of growth (i.e., before sporulation) in sporulation medium. Percent survival was calculated by comparing the treated CFU/mL (those exposed to 10mM H_2_O_2_) to untreated CFU/mL (water). While about 50% of the wild-type and the *dhb* mutant strain populations survived oxidative stress exposure, less than 1% of the *asb* mutant strain population survived (Figure 4D). This was due to a four-log decrease in the CFU/mL of the *asb* mutant strain following treatment with 10mM H_2_O_2_ (Figure 4C). The defect in survival was rescued by supplementation of the *asb* mutant strain with 25μM of purified petrobactin to the medium at the time of inoculation (Figure 4C, 4D). These data confirm our hypothesis that petrobactin is protective against oxidative stress during stationary phase—but prior to sporulation—in sporulation medium, which likely supports efficient sporulation and thus transmission between mammalian hosts.

### Petrobactin is preferred for rapid growth and sporulation in bovine blood

Following death of an infected mammal, blood laden with *B. anthracis* is exposed to the atmosphere by either hemorrhagic draining or the activity of scavengers on the carcass (11,44,45). Since vegetative bacilli are not easily infectious, *B. anthracis* transmission requires sporulation in aerated blood, a process triggered when the blood-borne CO_2_ reported to suppress sporulation decreases following death, thus triggering the sporulation cascade in a race against decomposition (44). Experiments to test *in vivo* sporulation are ethically and technically challenging, so to determine the relevance of each iron acquisition system—petrobactin, hemin, and bacillibactin—to disease we measured sporulation in bovine blood. Cultures of wild-type *B. anthracis* Sterne, the *asb* mutant, the *dhb* mutant, and the *isd* mutant strain (a mutant in hemin utilization) were grown in defibrinated bovine blood with shaking for three days. Every 24 hours, the total and sporulated CFU/mL were enumerated.

Compared to wild-type at 24 hours, growth of the *asb* mutant strain was reduced by one log (Figure 5A) with two log fewer spores (Figure 5B) whereas all other strains—the *isd* and the *dhb* mutant strains—had equivalent CFU/mL. While percent sporulation at 24 hours is low, generally < 25%, most sporulation in the wild-type strain appears to occur during the first 24 hours of incubation, after which non-sporulated cells begin to die thus reducing the total CFU/mL and increasing percent spores. Conversely, the *asb* mutant strain demonstrated delayed sporulation, gaining an additional log of spores between the 24 and 48-hour timepoints, but the percent sporulation at 48 hours and 72 hours was < 25% compared to the wild-type strain at 80% (Figure 5C). Percent sporulation for both the *isd* mutant strain and the *dhb* mutant strain were about 50%, though these were not statistically significant from the wild-type strain (Figure 5C). Additionally, both total and spores CFU/mL for both the *isd* and the *dhb* mutant strains were like wild-type, suggesting that petrobactin is a preferred iron gathering system during growth in bovine blood (Figure 5A, 5B).

The growth defect and delayed sporulation of the *asb* mutant strain could be due to oxidative stress, a lack of available iron, or a combination of the two stresses. To separate the effects of petrobactin supplementation on iron acquisition and protection from oxidative stress, the *asb* mutant strain was supplemented with 25μM of either petrobactin or hemin (n=2). Hemin is the oxidized form of heme, which is released into blood by the lysis of red blood cells and can be bound by *B. anthracis* hemophores, making it biologically relevant (46, 47). However, hemin isn’t known to protect against intracellular oxidative stress, so we predicted that if petrobactin were only required for iron acquisition, then hemin supplementation should complement the *asb* mutant strain phenotype.

Supplementation of the *asb* mutant with hemin did not affect overall growth but appeared to enhance early sporulation whereas supplementation with petrobactin rescued both growth and sporulation (Figure 5A-C). These data suggest that the iron provided via hemin may allow for efficient sporulation while the dual benefits of petrobactin iron acquisition plus protection from oxidative stress enable continued growth prior to the onset of sporulation.

**Figure 5.**
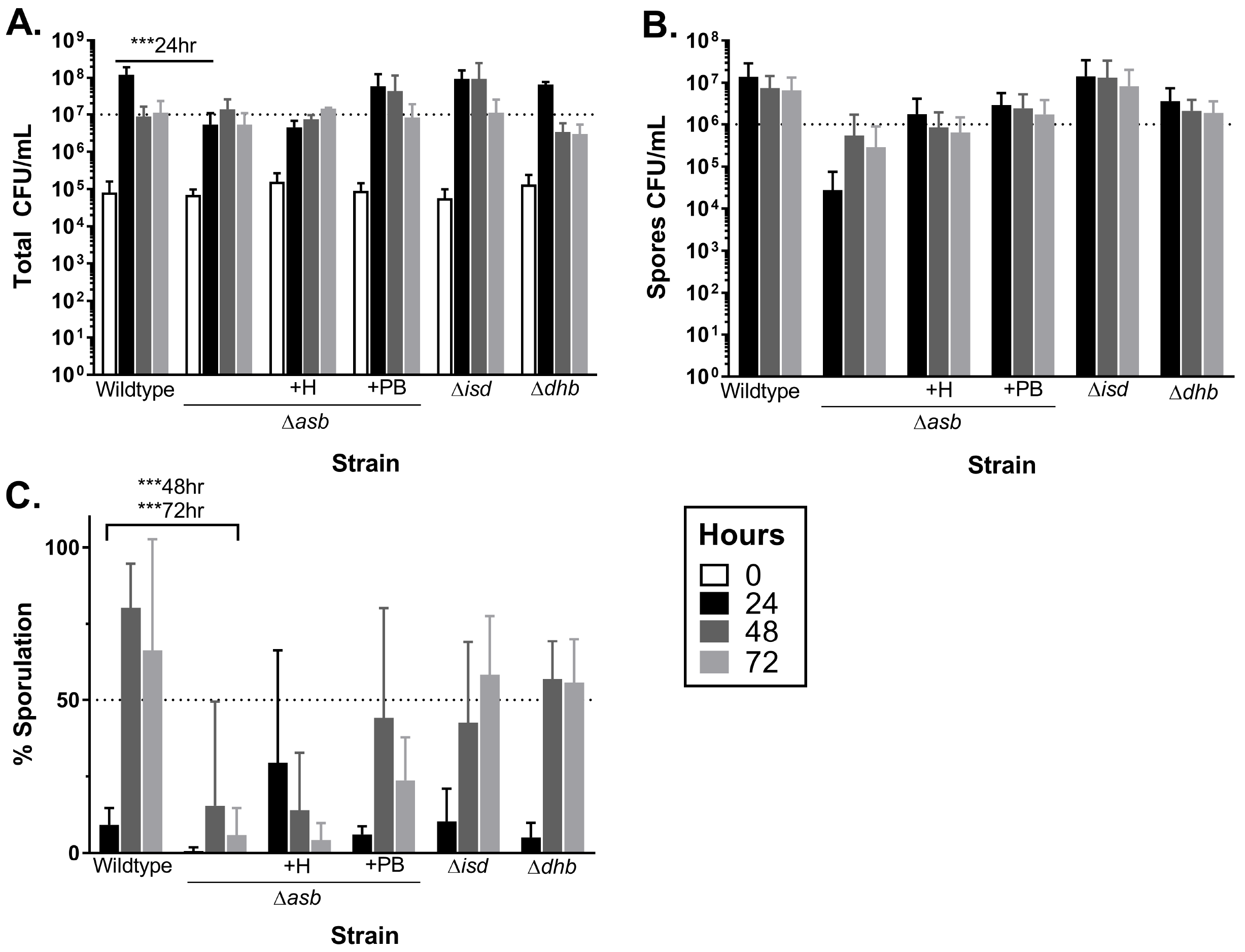
Petrobactin, but not hemin, is preferred for both growth and sporulation in bovine blood. Wild-type, the *asb* mutant, the *isd* mutant or the *dhb* mutant (supplemented with either 25μM of petrobactin (+PB) or 25μM hemin (+H) as indicated) were grown in defibrinated bovine blood. At 24 (black), 48 (dark gray), and 72 (light gray) hours post-inoculation, CFU/mL was determined for both A) total and B) spores and used to calculate the C) percent sporulation. Data are compiled from three independent experiments (except for *asb* mutant + 25μM hemin, n=2) and were analyzed using a two-way ANOVA with a Tukey’s multiple comparisons test ***p-value≤0.001. Dotted lines are placed to facilitate comparisons between strains and time points.

## Discussion

In this work, we show that petrobactin is not required for *B. anthracis* Sterne germination (Figure 1A) but is required for efficient sporulation in sporulation medium (Figure 1B). Using fluorescent *asbA:gfpmut3α* reporter fusions, we also show that *asb* is both transcribed and translated during late stage growth of *B. anthracis* Sterne prior to sporulation in a sporulation-sigma-factor independent manner (Figure 2B-C, 3). Unlike during vegetative growth, petrobactin is not exported during sporulation but remains intracellular (Figure 4A) where it has a significant role in protecting against oxidative stress (Figure 4C-D) and eventually associates with the spore (Figure 4B). These findings may have relevance to transmission since petrobactin is also required for efficient sporulation in bovine blood (Figure 5), a pre-requisite for survival and transmission of the pathogen (11, 44). We believe this to be the first demonstration that a siderophore is induced in preparation for sporulation and present in the mature spore.

The iron-gathering capacity of siderophores has long been appreciated for their role in pathogenicity and since their discovery, evidence for alternate functions has accumulated. Multiple reports have demonstrated roles for siderophores in: cell signaling, sporulation initiation, protection from copper and oxidative stress, the generation of oxidative stress against competitors, and, most recently, in survival via spores (48, 49).

In early 2017, Grandchamp et. al showed with *B. subtilis* that siderophore supplementation (including with the native bacillibactin) caused the onset of sporulation to occur earlier (49). Since this enhancement required import of the siderophore into the bacterial cell and iron removal by corresponding hydrolases, these authors hypothesized that the extra intracellular iron acted as a signal for the onset of sporulation (49). However, their study did not address bacillibactin regulation, export during sporulation, nor the cell stresses associated with
sporulation. So, to our knowledge, this is the first demonstration that a siderophore is induced to protect against oxidative stress prior to sporulation in high iron conditions.

As noted in the introduction, siderophores are primarily regulated by the iron-dependent repressor Fur. However, some siderophores, such as petrobactin, are biosynthesized in response to oxidative stress conditions and other catecholate-containing siderophores (*e.g.*, enterobactin and salmochelin) are protective against reactive oxygen species (35, 50–55). This protection is not due to iron sequestration that prevents additional Fenton reactions but is a function of the antioxidant properties of catechols (50, 53). Supplementation with free catechols doesn’t rescue the protective function of enterobactin, which requires import and hydrolysis for effective oxidative stress protection (53, 54). It is unclear whether petrobactin requires additional processing to become active against oxidative stress, though the detection of petrobactin associated with spores by mass spectrometry suggests it does not.

There are two non-exclusive hypotheses for siderophore upregulation during oxidative stress: one, that superoxide radicals oxidize iron co-factors thus inactivating key enzymes and two, that upregulation of the enzymes to mitigate oxidative stress require metallic (*e.g.*, iron and manganese) co-factors. Both of these would reduce the intracellular iron pool and thereby relieve iron from Fur to enable iron acquisition system expression (35, 51). In the case of *Bacillus* spp., the intracellular iron pool is further depleted during the onset of sporulation due to upregulation of aconitase, an iron-rich citrate isomerase and stabilizer of σ^K^-dependent gene transcripts (56, 57).

While the demand for iron during oxidative stress and/or sporulation may relieve negative regulation by Fur, it is likely that *asbA-F* expression is induced by an oxidative stress regulator such as PerR. Enterobactin is positively regulated by the oxidative stress response and there is compelling evidence linking *Azotobacter vinelandii* catecholate siderophores to similar regulation (53–55). The observed phenotype for those siderophores is similar to that observed by Lee et al. for petrobactin: high iron repression of the siderophore can be overcome by oxidative stress (35,53,55). Petrobactin biosynthesized within the cell may then become randomly associated with the prespore. More work is needed to better characterize regulation of the *asb* operon and petrobactin biosynthesis.

As growth in blood marks an endpoint for an anthrax infection, the bacilli must not only grow well, but also prepare for survival and transmission between hosts. Evidence in the literature suggests that exposure of blood-borne bacilli to oxygen as a dying host bleeds out begins the signaling cascade for sporulation creating a direct link between growth and sporulation in blood and transmission (11, 44). It’s known that petrobactin is required for growth in macrophages and iron-depleted medium, but that requirement had not been demonstrated for growth or sporulation in blood prior to these experiments. Our data suggest that petrobactin is the preferred iron acquisition system for growth and sporulation in bovine blood, despite multiple potential iron sources. While, petrobactin was required to achieve wild-type growth of 10^8^ CFU/mL in blood, the *asb* mutant was still able to grow to 10^7^ CFU/mL suggesting that another iron acquisition source was functioning, likely either the *isd* system or bacillibactin. More work is needed to fully understand the contributions of each iron system to growth and sporulation and to verify these findings in other *B. anthracis* strains.

These data update the model of *B. anthracis* Sterne iron acquisition and sporulation (Figure 6). In this model, upon entry of the bacterial population into late stage growth, environmental stressors both deplete the intracellular iron pool and induce oxidative stress that act to upregulate the *asb* operon, presumably through PerR regulation. Petrobactin is biosynthesized for iron acquisition and/or protection against oxidative stress, which support the bacillus as it transitions into sporulation. Either direct import of petrobactin into the prespore or random association results in packaging of petrobactin into the spore. These findings underscore the vital role of petrobactin in the many stages of *B. anthracis* infection, from survival in the macrophage to growth in the bloodstream and now, sporulation, which facilitates transmission to a new host.

**Figure 6.**
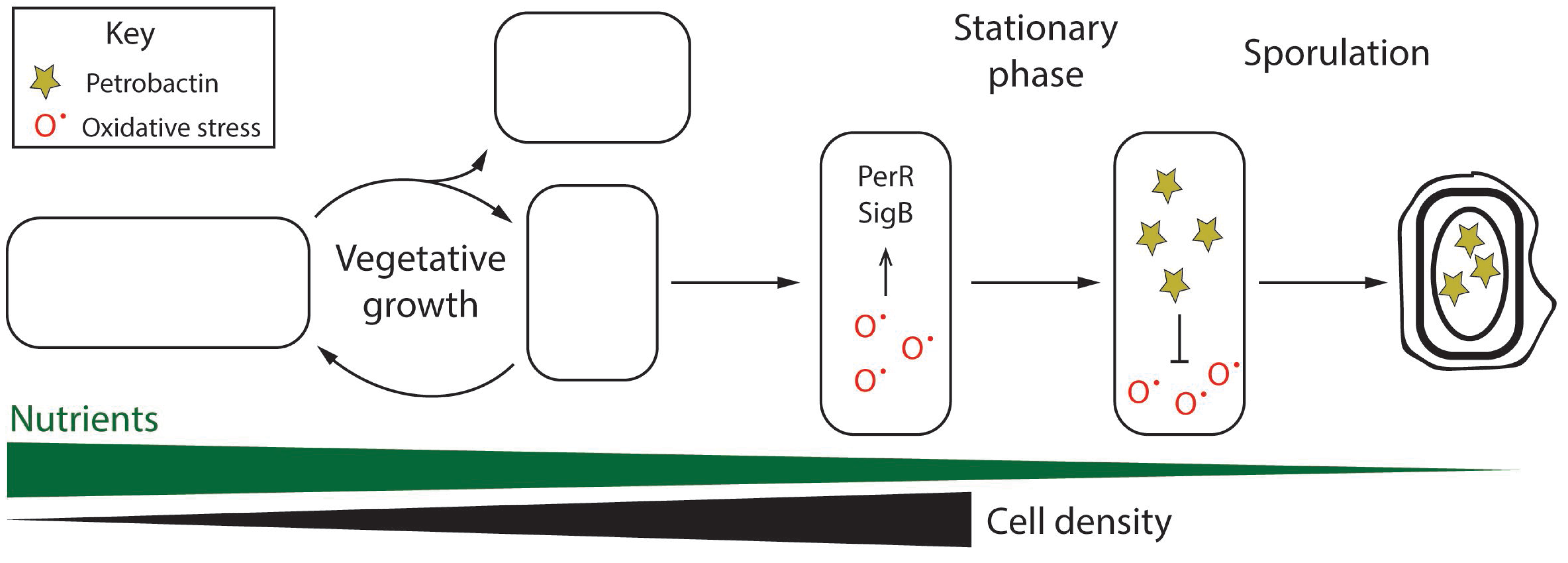
Proposed model of petrobactin use by *B. anthracis* during late stage growth and early sporulation.

## Materials and Methods

### Bacterial growth conditions and sporulation

Strains used are described in Supplementary Table 1. Genomic-based fluorescent reporters were generated by PCR amplification and Gibson cloning (New England Biolabs) of the genetic construct into pBKJ258 which was then inserted onto the *Bacillus anthracis* Sterne 34F2 (pXO1^+^ pXO2^−^) genome by allelic exchange, as described by Janes and Stibitz (58). The plasmid-based transcriptional reporter was directionally cloned with EcoRI and BamHI into the pAD123 multiple cloning site upstream of the promoterless *gfpmut3a*. All necessary primers are listed in Supplementary Table 2. Modified G medium (ModG) was used for the generation of *B. anthracis* spores at 37°C for 72 hours (59). Spores were collected at 2,800 rpm then washed and stored in sterile water at room temperature following heat activation at 65°C. Strains containing plasmid-based reporters (39) were grown in the presence of 10μg/mL chloramphenicol. Media and chemicals were purchased from Fisher Scientific or Sigma Aldrich.

### Spore germination

Spore germination was measured in iron-depleted medium (IDM) supplemented with 1mM inosine, following a 20 minute heat activation at 65°C (29). To measure germination and subsequent outgrowth, spores were inoculated at a starting OD_600_ between 0.25 and 0.5 for a final volume of 200μL (n=3). The spores were incubated at 37°C in a SpectraMAX M2 spectrophotometer and the OD_600_ measured every five minutes for one hour. Data are representative of three independent experiments and are presented as percent of the initial OD_600_.

### Supplementation of *asb* mutant sporulation with petrobactin and outgrowth

To supplement *asb* mutant spores with petrobactin, bacilli were grown overnight at 30°C in BHI (Difco) inoculated 1:1000 in 25mL of ModG medium supplemented with 25μM of purified petrobactin. After a 72-hour incubation, spores were collected by centrifugation at 2,800rpm and washed three times with 20mL of sterile, deionized water. The spores were resuspended in 1mL of water following heat activation for 20 minutes at 65°C.

### Reporter growth, measurement, and analysis

Bacterial strains were plated on BHI and grown in BHI at 30°C overnight. Overnight cultures were back diluted 1:50 into fresh BHI and incubated at 37°C for one hour. The cells were pelleted at 2,800 rpm for 10 minutes (Centrifuge 5810 R, Eppendorf), washed once with BHI then used to inoculate 200μL of ModG medium. Each strain was inoculated into triplicate wells of a 96-well plate to a starting OD_600_ of 0.05, covered with a gas permeable sealing membrane (Breathe-Easy, Diversified Biotech) then grown in a Synergy HTX plate reader at 37°C with continuous shaking at 237cpm for 12 hours. The OD_600_ and fluorescence (excitation 485/20, emission 528/20) were bottom-read every five minutes using a tungsten light source. Data were analyzed in R software by first subtracting a media blank from both fluorescence and OD_600_ then normalizing fluorescence by the OD_600_ (60). Background fluorescence was approximated by wild-type cells and subtracted from the reporters at corresponding timepoints.

### Microscopy

Wild-type and the *asb* translational reporter expressing Gfpmut3α were grown in ModG medium at 37°C and every two hours from six to 12 hours post-inoculation, five μL was spotted on a microscope slide. At least 100 bacteria were imaged at each timepoint scored for fluorescence and/or sporulation. Bacteria were counted by determining the size of a bacterium and calibrating all images to this length. A bacterium was scored as positive for fluorescence if the intensity was at least 1.4× above the background fluorescence of wild-type, non-Gfpmut3α-expressing bacilli. A bacterium was positive for sporulation upon observation of a phase bright spore. Phase-contrast and fluorescence microscopic images were taken using a Nikon TE300 inverted microscope equipped with a mercury arc lamp, 60× Plan-Apochromat 1.4-numerical aperture objective, cooled digital CCD camera (Quantix Photometrics). Excitation and emission wavelengths were selected using a 69002 set (Chroma Technology) and a Lambda 10-2 filter wheel controller. Fluorescence images of Gfpmut3α were captured with excitation and emission filters centered at 490nm and 535nm, respectively. Exposures was set at 300ms.

### Oxidative stress survival

Wild-type, *asb* mutant, and *dhb* mutant strains were grown in ModG medium. At eight hours post-inoculation, 500μL of each culture was added to 100μL of either sterile water or 60mM hydrogen peroxide (final concentration 10mM). Treated (10mM H_2_O_2_) and control (H_2_O) cultures were incubated for 10 minutes at 37°C then serially diluted in PBS and plated on BHI to stop the reaction and count CFU/mL. Any culture below the limit of detection (about 667 CFU/mL), were assigned a conservative value of 600 for data analysis. Data are pooled from three independent experiments and presented as percent survival: treated divided by untreated times 100.

### LAESI-MS

Samples from ModG medium for LAESI-MS were collected at 12 hours postinoculation and separated by centrifugation to obtain the culture medium and cell pellets. Cells were washed once in an equal volume of PBS. All samples were stored at -80°C until analysis. Spores for LAESI-MS analysis were prepared as described above. Unless indicated otherwise, 6×10^7^ spores from three independent spore preparations in 50% DMSO (or 15μL of cell pellets) were plated in triplicate wells of shallow 96-well plates and subjected to laser-based ablation.

The ESI mass spectrograph was obtained using a ThermoFisher LTQ XL mass spectrometer, containing an atmospheric pressure ionization stack with a tube lens and skimmer, three multipoles, a single linear trap configuration and a set of 2 electron multipliers with conversion dynodes. The mass spectrometer was connected to a Protea LAESI DP-1000 instrument with an ESI electrospray emitter for ambient ionization. The collected data points were exported to Gubbs™ Mass Spec Utilities (61) and processed using Generic Chromatographic Viewer for individual *m/z* (ThermoFisher Scientific). The average intensity of petrobactin was normalized as necessary (e.g., OD_600_ or 10^6^ spores).

### Sporulation efficiency

Wild-type, *asb, dhb,* and *isd* mutant strains were grown in BHI either overnight at 30°C then inoculated at 1:1000 into three mL of either ModG medium or defibrinated bovine blood (Hemostat laboratories). Cultures were grown at 37°C with growth and sporulation enumerated at regular intervals by serial dilution in PBS prior to plating on BHI for growth at 37°C overnight. Cultures were plated both before and after a heat treatment step (30 minutes at 65°C) to obtain total and spores, respectively. CFU/mL below the limit of detection (~667) were assigned a conservative value of 600 for data analysis. Percent sporulation is post-heat-treatment divided by the total times 100. Hemin for supplementation was first suspended at 3.83mM in 1.4M NaOH, then diluted to 150μM in PBS (62). Data are pooled from three independent experiments unless otherwise noted.

## Author contributions

A.K.H. was responsible for project and experiment design, data analysis, spore harvests, construct design, blood sporulation experiments, and drafting the manuscript. Y.P. completed ModG sporulation and oxidative stress experiments. R.D. and S.C. constructed reporter plasmids and strains. Z.M. performed microscopy and image processing. A.T. processed and analyzed petrobactin content by LAESI-MS. D.S., A.T., and P.C.H. provided funding, resources, and conceptual advice. All authors contributed to the final manuscript. No authors report a conflict of interest.

## Funding

Funding for this work was provided by the NIH (R35 GM118101, D.H.S.; T32 AI00758, A.K.H.), the UM-Israel Partnership for Research (P.C.H.), the UM Endowment for the Basic Sciences Innovation Initiative (P.C.H., D.H.S, A.T., A.K.H.), UM Rackham Graduate School (A.K.H.), the American Society for Microbiology Watkins Fellowship (A.K.H.), and the Hans W. Vahlteich Professorship (D.H.S.). The sponsors had no role in study design, data collection and interpretation, or the decision to submit the work for publication

## Acknowledgements

Authors would like to thank Dr. Suzanne Dawid for her thoughtful suggestions regarding manuscript structure and acknowledge Nick Lesniak and Dr. Marc Sze for their comments. This manuscript was conditionally read by the U.S. Department of Homeland Security upon acceptance.

---

**Supplementary Table 1. Strains of *B. anthracis* Sterne 34F2 used in this work.**

**Supplementary Table 2. Primers used to generate mutant strains used in this work.**

**Supplementary Figure 7. Phase contrast and fluorescent imaging of wild-type *B. anthracis* Sterne bacilli during growth in sporulation medium.** The strain was grown in ModG sporulation medium with both phase-contrast and fluorescent (excitation: 490nm and emission: 535nm) micrographs taken at six, eight, ten, and 12 hours of growth. Representative images from each time point. **A)** six hours **B)** eight hours **C)** ten hours **D)** 12 hours.

